# Chr21 protein-protein interactions: enrichment in products involved in intellectual disabilities, autism and Late Onset Alzheimer Disease

**DOI:** 10.1101/2019.12.11.872606

**Authors:** Julia Viard, Yann Loe-Mie, Rachel Daudin, Malik Khelfaoui, Christine Plancon, Anne Boland, Francisco Tejedor, Richard L. Huganir, Eunjoon Kim, Makoto Kinoshita, Guofa Liu, Volker Haucke, Thomas Moncion, Eugene Yu, Valérie Hindie, Henri Bléhaut, Clotilde Mircher, Yann Herault, Jean-François Deleuze, Jean-Christophe Rain, Michel Simonneau, Aude-Marie Lepagnol-Bestel

## Abstract

Intellectual disability (ID) found in Down syndrome (DS), which is characterized by an extra copy of 234 genes on Chr21 is poorly understood. We first used two DS mouse models that either display an extra copy of the *Dyrk1A* gene or of the mouse Chr16 syntenic region. Exome sequencing of transcripts deregulated in embryonic hippocampus uncovers enrichment in genes involved in chromatin and synapse respectively. Using large-scale yeast two-hybrid screen (154 distinct screens) of human brain library containing at least 10^7^ independent fragments, we identified 3,636 novel protein-protein interactions with an enrichment of direct interactors of both Chromosome 21(Hsa21) baits and rebounds in ID-related genes. Using proximity ligation assays, we identified that Hsa21-encoded proteins are located at the dendritic spine postsynaptic density in a protein network located at the dendritic spine post synapse. We located Hsa21 DYRK1A and DSCAM that confers a ∼ 20-fold increase in Autism Spectrum Disorders (ASDs), in this postsynaptic network. We found that a DSCAM intracellular domain binds either DYRK1A or DLGs that are multimeric scaffolds for the clustering of receptors, ion channels, and associated signaling proteins. The DYRK1A-DSCAM interaction is conserved from drosophila to humans. The identified postsynaptic.network is enriched in ARC-related synaptic plasticity, ASDs and Late-Onset Alzheimer Disease. Altogether, these results emphasize links between DS and brain diseases with complex genetics.

## INTRODUCTION

Down syndrome (DS) is the most common form of intellectual disability (ID), with an incidence of one in 800 births. It is a human genetic disorder, caused by the presence of a third copy of 234 genes from *Homo sapiens* autosome 21 (Hsa21) (Antonarakis, 2017; Antonarakis et al., 2004). DS is one of the most complex viable genetic perturbations. In spite of a broad spectrum of clinical symptoms, features common to all DS variants are the intellectual deficit that impairs learning and memory and an increased risk of developing a dementia resembling Alzheimer’s disease (AD), even in patients as young as 40 years old (Ballard et al., 2016; Dierssen, 2012; Wiseman et al., 2015a). To date, the precise contribution of each Hsa21 protein overexpression to the cognitive impairment found in DS remains undetermined.

Here, we used two DS mouse models. The first model is a *Dyrk1A* BAC 189N3 model carrying a triplication of the ∼152 kb mouse *Dyrk1a* locus containing the whole mouse Dyrk1a gene with a 6 kb flanking fragment on the 5’ side and a 19 kb flanking fragment on the 3’ side (Guedj et al., 2012). The second model is a transgenic line (Dp(16)1Yey) carrying a triplication of 22.9 Mb from *Mus musculus* chr16 (Mmu16), which are syntenic to 119 genes from Hsa21, including *DYRK1A* (dual-specificity tyrosine phosphorylated and regulated kinase 1A) (Yu et al., 2010), thus precisely reflecting the gene dosage of Hsa21 orthologs. The *Dyrk1A* gene was found to be a major player in DS whom overexpression induces modifications in synaptic plasticity both in the hippocampus and in the prefrontal cortex (Ahn et al., 2006; Thomazeau et al., 2014). Dyrk1a is an important candidate suggested to be involved in the learning and memory impairment seen in DS patients (Smith et al., 1997), but the regulatory pathways impaired by trisomy of *DYRK1A* remain elusive.

In this work, we examined the respective contributions of *Dyrk1a* and other HSA21 gene products in pathways linked to ID. Exome sequencing of transcripts mis-regulated in the embryonic hippocampus uncover two contrasting repertoires of genes: chromatin related genes for the *Dyrk1A* trisomy model and a synapse related genes for the Dp(16)1Yey model. In order to gain further insight into the molecular network of proteins responsible for DS phenotypes, we next searched for human brain proteins that interact with proteins encoded by HSA21. To this end, we conducted a large-scale yeast two-hybrid screen using HSA21 baits and a human brain library of targets. This analysis revealed that both direct interactors of HSA21-encoded proteins and their direct rebounds are enriched in proteins involved in ID. Moreover, we found an enrichment in HSA21-encoded proteins that are part of a network located in dendritic spine postsynaptic density. The same interactome is also enriched with proteins involved in ARC-related synaptic plasticity, Autism Spectrum Disorders and Late-Onset Alzheimer Disease (LOAD).

## RESULTS

### Whole-genome RNA sequencing and quantitative proteomics reveal two contrasted networks of deregulated genes of hippocampus from 189N3 *DYRK1A* and Dp(16)1Yey DS models

We used the 189N3 and the Dp(16)1Yey/+ DS mouse models. We performed RNA sequencing on embryonic E17 hippocampi of these two DS models. We identified 84 deregulated genes in 189N3 mice (**Supplementary Table S1**) and 142 deregulated genes in Dp(16)1Yey/+ (**Supplementary Table S2**) compared to their littermate controls.

Gene ontology (GO) Process analyses of differentially expressed genes revealed a deregulation of chromatin proteins for 189N3 mice with:

> GO:0006334∼nucleosome assembly (P value=1.17 e^-08^)

> GO:0031497∼chromatin assembly (P value= 5.98 e^-08^)

> GO:0034728∼ nucleosome organization (P value = 1.71 e^-07^).

Deregulation of chromatin proteins is in agreement with data that previously reported by us (Lepagnol-Bestel et al., 2009; Loe-Mie et al., 2010).

In contrast, for Dp(16)1Yey/+, we found a deregulation of proteins involved in synaptic function:

> GO:0007268∼chemical synaptic transmission (Pvalue= 6.87e-^09^)

> GO:0051932∼synaptic transmission, GABAergic (Pvalue= 1.27e-

> 05)GO:0048812∼neuron projection morphogenesis (Pvalue= 8.24e-^05^)

Interestingly, for the GO:0051932∼synaptic transmission, GABAergic, 6 of 77 genes are deregulated for a repertoire of 53 out of 24850 human genes (7.8%), indicating an enrichment of 36.54 fold compared to expectations (hypergeometric P-value = 1.48 e-^08^). This result is in full agreement with the impairment of excitation-inhibition balance (E–I balance) in synaptic activity found in DS (Kleschevnikov et al., 2012; Raveau et al., 2018).

First, we used Dapple analysis (Rossin et al., 2011) of 189N3 and Dp(16)1Yey/+ differential expressed genes to evidence two contrasted networks with statistically significance (direct edge counts; *p*<0.05) showing that genetic variation induced by the triplication of the mmu16 affects a limited set of underlying mechanisms (**Figure 1 A-B**). We next used Webgestalt. Suite (Liao et al., 2019) for the142 deregulated genes in Dp(16)1Yey/+ with 77 upregulated and we identified two significant networks (A) and (B). **A**: a network that includes 8 genes from the 70 upregulated list with an enrichment in GO Biological Process: chemical synaptic transmission (p=220446e-16). **B**: a network that includes 4 genes from the 77 upregulated list with an enrichment in GO Biological Process: Biological Process: glutamate receptor signaling pathway (p=220446e-16). Note that three genes (*Camk2a*, *Gda*, *Dlgap3*) are part of a protein network of ARC-dependent DLG4 interactors that include 20 proteins (Fernández et al., 2017), indicating an over enrichment of 43.85 fold compared to expectations (hypergeometric p-value = 4.20 e-05) [Parameters: 3, 20, 77, 22,508 number of mouse genes from Mouse Ensembl (GRCm38.p6)].

**Figure 1.**
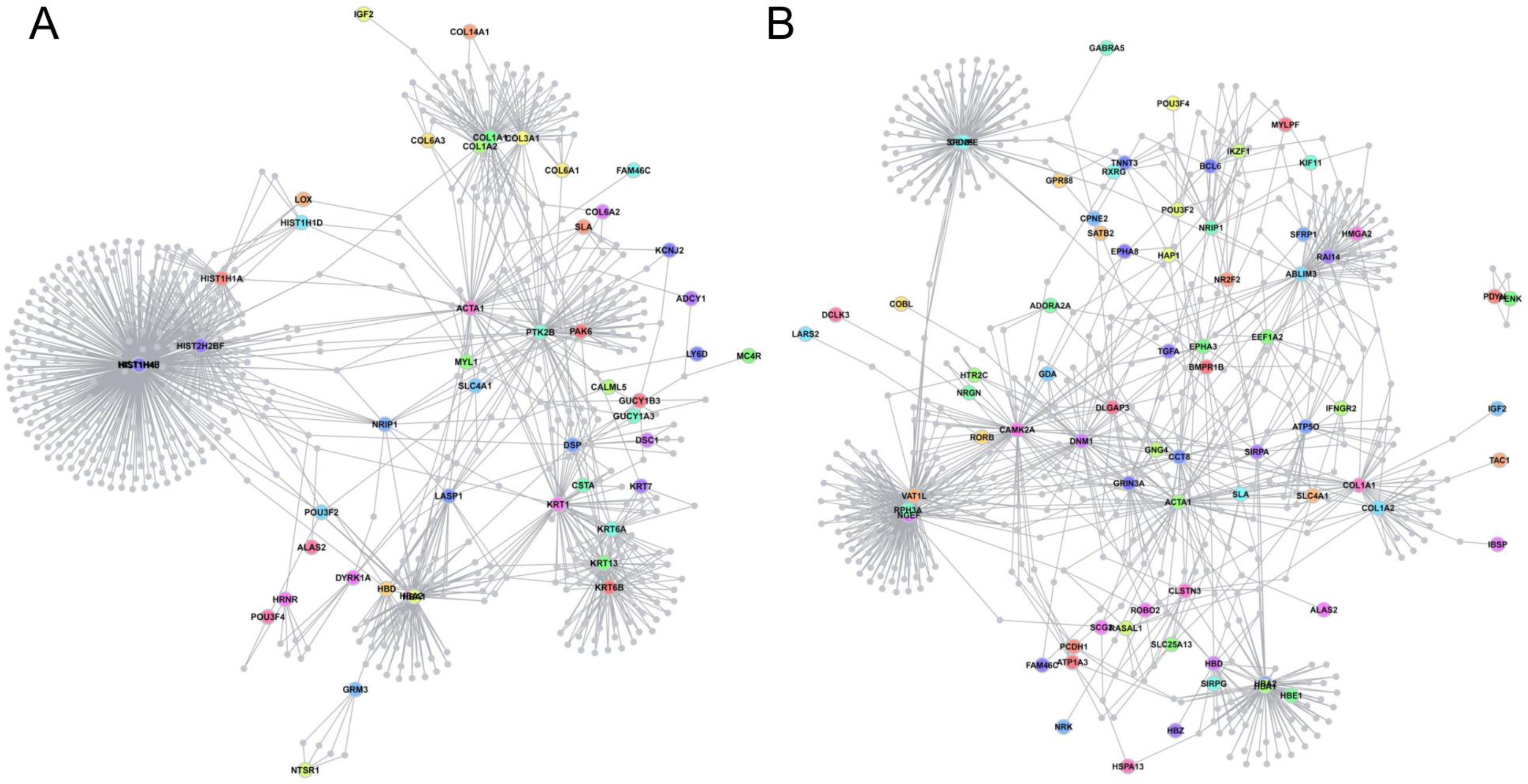
Protein-Protein Interaction Network generated from proteins encoded by deregulated genes identified in E17 hippocampus of 189N3 and Dp(16)1Yey transgenic mouse models, respectively. We performed RNA sequencing on embryonic E17 hippocampi of these two OS models. We identified 84 deregulated genes in 189N3 **(Supplementary Table Sl)** and 142 deregulated genes in Dp(16)1Yey/+ **(Supplementary Table S2)** compared to their littermate controls. They were included as an input to DAPPLE. Disease Association Protein-Protein Link Evaluator (DAPPLE), which uses high-confidence pairwise protein interactions and tissue-specific expression data to reconstruct a PPI network (Rossin et al., 2012).The network is conservative, requiring that interacting proteins be known to be coexpressed in a given tissue. Proteins encoded by deregulated genes are represented as nodes connected by an edge if there is in vitro evidence for high-confidence interaction. **A:** for 189N3 mice,we found an enrichment in Gene ontology (GO) Process analyses of differentially expressed genes revealed a deregulation of chromatin proteins for 189N3 mice with : GO:0006334∼nucleosome assembly (P value=1.17 e^-08^). The DAPPLE network based on the analysis of 75 genes is statistically significant for direct and indirect connectivity more than would be expected by chance. For direct connectivity: Direct edges Count 26;expected 5.5; permuted 9.99 x 10^-4^; Seed Direct Degrees Mean 2.26: expected 1.27; permuted p=9.99 x 10^-3^. For indirect connectivity, Seed Indirect Degrees Mean 52.54; expected 29.43;permuted 9.99 x 10^-4^; Cl Degrees Mean 2.51;expected 2.28; permuted p=2.99 x 10^-3^. **B:** for Dp(16)1Yey/+,we found a deregulation of proteins involved in synaptic function :GO:0007268∼chemical synaptic transmission (Pvalue= 6.87e^-09^). The network based on the analysis of 131 genes is statistically significant for direct connectivity more than would be expected by chance (Direct edges count, 16;expected 7.933;permuted p=1.09 x 10^-2^). genes

**Figure 2.**
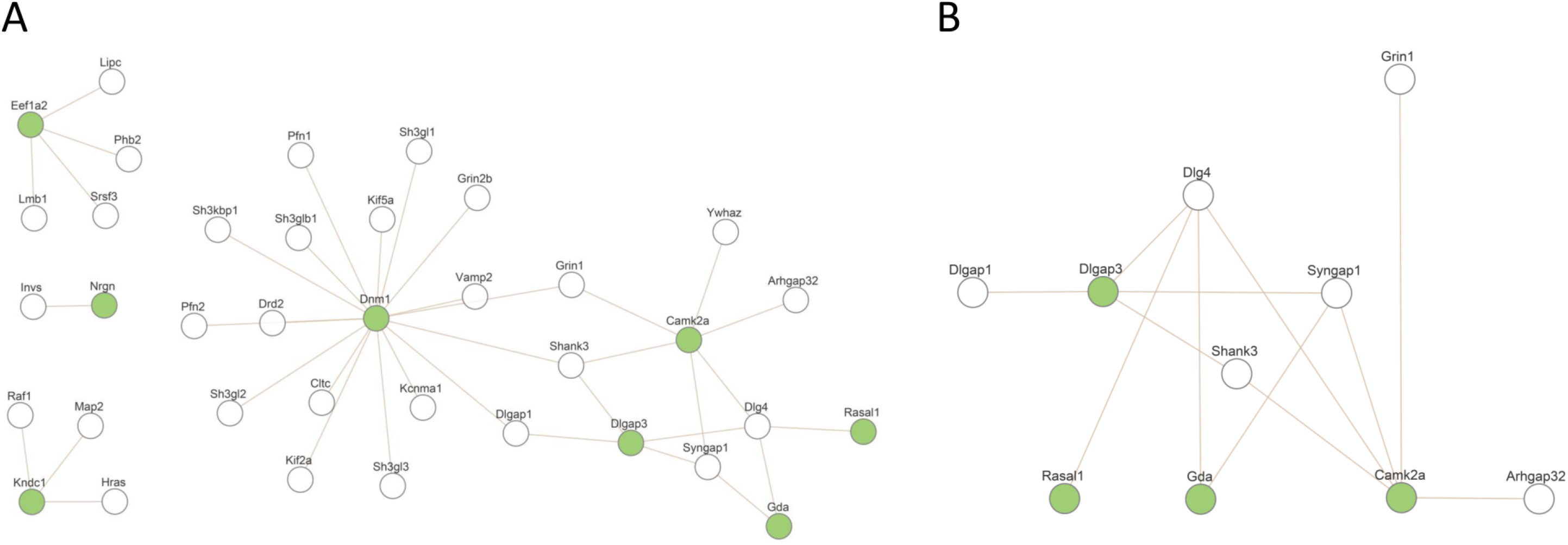
Products of deregulated genes in Dp(16)1Yey/+ are enriched in proteins linked to glutamate receptor signaling pathway and in proteins involved in an ARC-PSD95 complex linked to ID and intelligence. We found 142 deregulated genes in Dp(16)1Yey/+ with 77 upregulated. Using Webgestalt suite,we identified two significant networks (A) and (B). **A:** a network that includes 8 genes from the 70 upregulated list with an enrichment in GO Biological Process: chemical synaptic transmission (p=220446e-16). **B:** a network that includes 4 genes from the 77 upregulated list with an enrichment in GO Biological Process: Biological Process: glutamate receptor signaling pathway (p=220446e-16). Note that the 3 genes (Camk2a, Gda, Dlgap3) are part of a protein network of ARC-dependent DLG4 interactors that include 20 proteins (Hernandez et al., 2018), indicating an over enrichment of 43.85 fold compared to expectations (hypergeometric p-value = 4.20 e-05). [Parameters: 3, 20, 77, 22,508 number of mouse genes from Mouse Ensembl (GRCm38.p6)].

Altogether, these results indicate a contrasted deregulation of chromatin-related genes for 189N3 model and synaptic plasticity-related genes with a n enrichment in genes linked to ARC postsynapse Complexes Involved with Neural Dysfunction and Intelligence for Dp(16)1Yey/+ model.

### Establishment of a Hsa21 protein-protein interaction map by a high-throughput, protein domain-based yeast two-hybrid (Y2H) screening

To improve our knowledge of the molecular network underlying DS, we performed a large-scale protein-protein interaction study. We performed 72 screens with HSA21 protein as baits and 82 screens against their direct interactors (rebounds) using a highly complex random-primed human adult brain cDNA library. We identified 1687 and 1949 novel direct interactions (**Figure 3 A-D**). These interactions were ranked by category (a to f), using a Predicted Biological Score (PBS) (Formstecher et al., 2005). Analysis of direct interactors from 72 HSA21 baits screens gives 1687 novel interactions identified with 76 already known (Biogrid) interactions confirmed (**Figure 3A-B**). Analysis of direct interactors from 82 rebound screens gives 1949 novel interactions were identified with 97 already known (Biogrid) interactions confirmed (**Figure 3 C-D**). We next compared these direct interactors with three lists of genes involved in intellectual disability (ID) (Deciphering Developmental Disorders Study, 2015; Gilissen et al., 2014). We generated three datasets of ID genes: S10 list (n=527), S11 list (n=628) and S2 (n=1244) (Supplementary Table S3). HSA21 direct interactors are enriched in ID proteins (X21 baits against S10: p-value = 2.29e-08; X21 baits against S11: p-value = 9.39e- 12; X21 baits against S2: p-value = 7.53e-13) (**Figure 3 E**). Similarly, HSA21 rebound direct interactors are enriched in ID proteins (X21 rebounds against S10: p-value = 8.30e-09; X21 rebounds against S11: p-value = 8.64e-12; X21 rebounds against S2: p-value = 7.76e-14) (**Figure 3 F**). Altogether, these results indicate that both HSA21 direct interactors and rebound direct interactors are part of a large ID network.

**Fig. 3.**
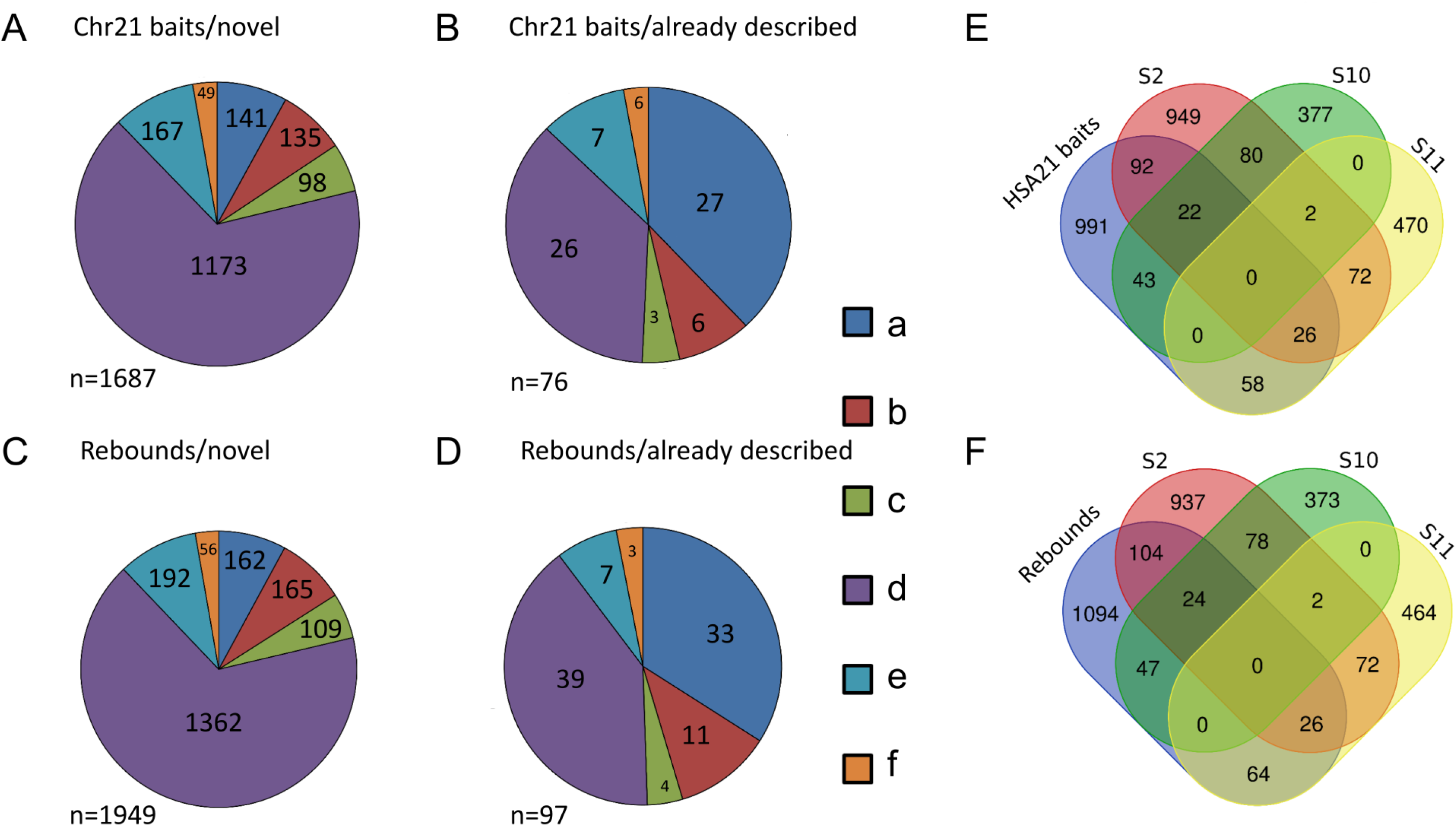
high-throughput Y2H identifies 3636 novel direct interactions with their enrichment in proteins involved in Intellectual Disabilities. 72 screens with HSA21 protein as baits and 82 screens against their direct interactors (rebounds) have been performed using a human brain library. 1687 and 1949 novel direct interact ions have been identified. These interactions were ranked by category (a to f), using a Predicted Biological Score (PBS) (Formstecher et al.,2005). Analysis of direct interactors from 72 HSA21 baits screens **(A-C)**. 1687 novel interactions were identified **(A)** and 76 already known (Biogrid) interactions confirmed **(B).** Analysis of direct interactors from 82 rebound screens **( D-F).** 1949 novel interactions were identified **(D)** and 97 already known (Biogrid) interactions confirmed **(E).** We compared these direct interactors with three lists of genes involved in Intellectual Disability (Gilissen et al. Nature, 2014; Deciphering Developmental Disorders Study,2015). Both HSA21 direct interactors (**C**) and rebound direct interactors **(F)** are enriched in ID proteins (see text) suggesting that these two types of interactors are part of a large ID network.

We realized a biological processes analysis using GO DAVID (see methods) (**Figure 4**). The colored nodes correspond to the most significant results with GO:0022008∼Neurogenesis (*p*- value=3.06e-17); GO:0048812∼Neuron projection morphogenesis (*p*-value=, 2.91e-13); GO:0050767∼Regulation of neurogenesis (*p*-value=2.66e-06); GO:0043632∼Modification-dependent macromolecule catabolic process (*p*-value=6.46e-05); GO:0051962∼Positive regulation of nervous system development (*p*-value=6.29e-06); GO:0045665∼Negative regulation of neuron differentiation (*p*-value=1.55e-07). Altogether, our data indicate an enrichment of interactions related to neuronal differentiation.

**Figure 4:**
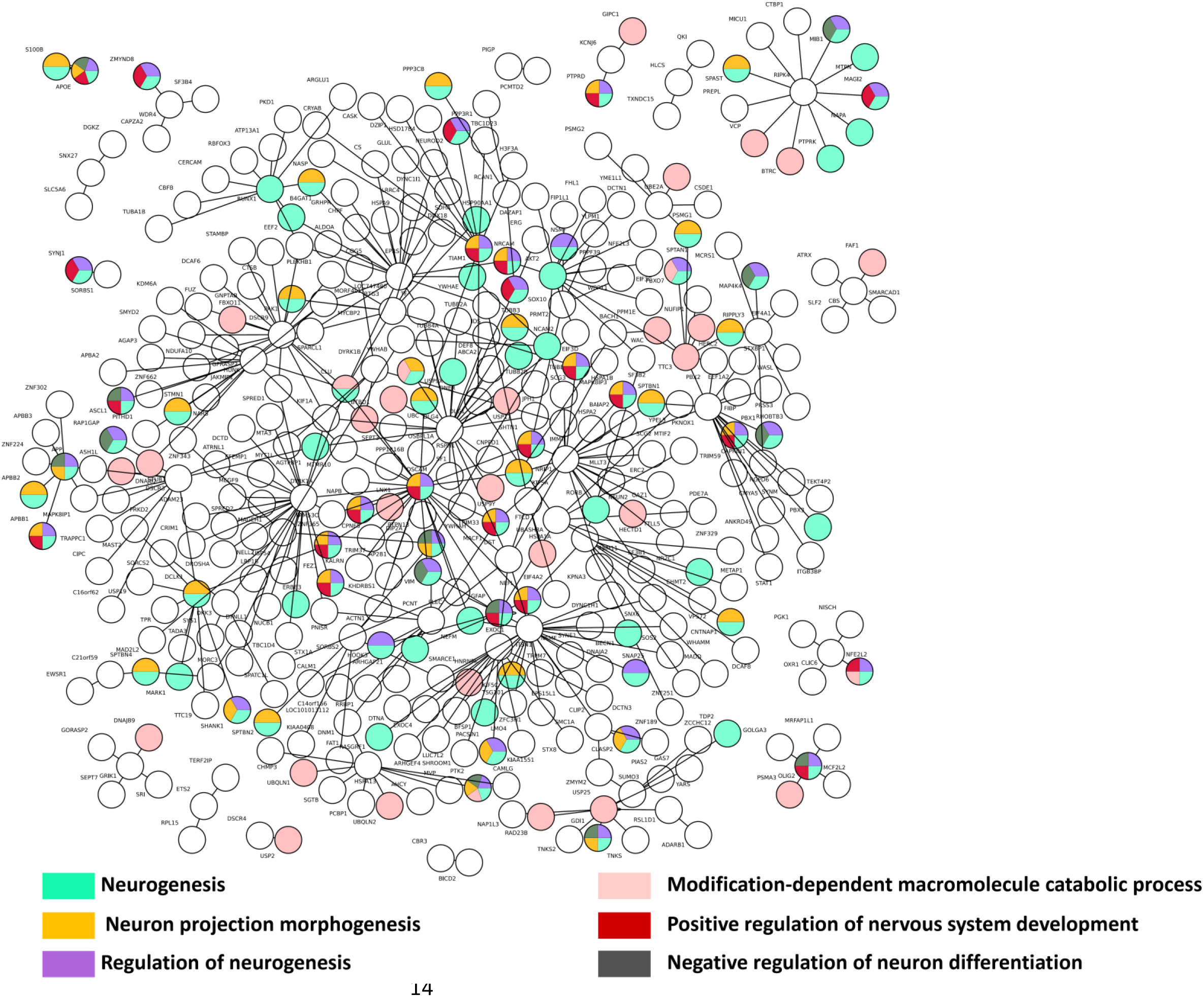
Biological processes network interactions from Yeast two-hybrid protein-protein interaction data. A biological processes analysis using GO DAVID was realized (see methods). The colored nodes correspond to the most significative results: GO:0022008∼Neurogenesis; GO:0048812∼Neuron projection morphogenesis; GO:0050767∼Regulation of neurogenesis; GO:0043632∼Modification-dependent macromolecule catabolic process; GO:0051962∼Positive regulation of nervous system development; GO:0045665∼Negative regulation of neuron differentiation with *p*-value 3.06e-17, 2.91e-13, 2.66e-06, 6.46e-05, 6.29e-06, 1.55e-07, respectively). A color correspond to a cluster of several biological processes. The multi-colored nodes correspond to genes presents in different annotation clusters.

### Linking HSA21 proteins to Late Onset Alzheimer Disease (LOAD) and neuropsychiatric diseases: DSCR9-CLU, DYRK1A-RNASEN

We first focused our analysis to two novel interactions with a potential importance in brain diseases, by combining Yeast two Hybrid (Y2H) interaction data with proximity ligation assays (PLA) that allow to localize PPI at the subcellular level when the maximal distance between the antibodies required for producing a signal is 40 nm (Söderberg et al., 2006).

The two selected PPI links respectively HSA21 DSCR9 gene with CLU, a risk factor for Late-Onset Alzheimer Disease (LOAD), HSA21 DYRK1A with RNASEN, a microprocessor complex subunit (**Figure 5A**).

**Figure 5.**
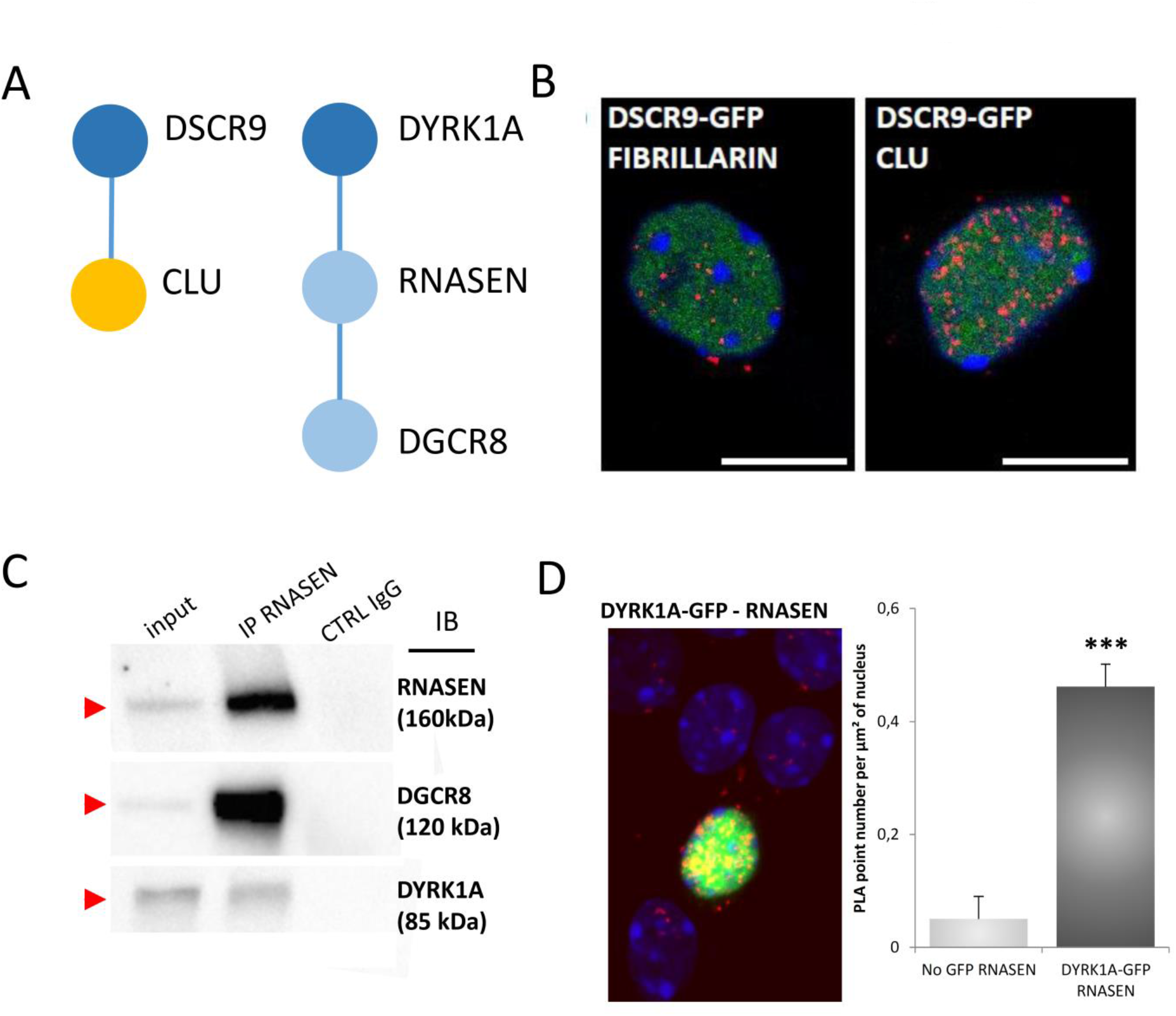
Interactions of HSA21 proteins with proteins involved in LOAD, Intellectual Disability and neuropsychiatric diseases. **(A).** Schematic representation of protein-protein interactions identified by yeast-two-hybrid using a human brain library. Dark blue circles indicate HSA21 encoded proteins; orange circle indicates a LOAD-related protein. **(B).** *In situ* PLA on primary cortical neurons transfected at DIC5 and fixed 48 hours later at DIC7 (red fluorescence) using anti-GFP and anti-Clu antibodies. PLA using anti-GFP and anti-Fibrillarin antibodies were performed as a negative control. Green fluorescent protein was visualized on green channel and nuclear bodies were labelled using Topro3 (blue fluorescence) . **(C)** and **(D):** DYRK1A interaction with RNASEN. **(C):** EK293 cells were immunoprecipitated (IP) using anti-RNASEN antibody and anti-lgG antibody as a negative control. The input and precipitated fractions were then resolved by sodium dodecyl sulphate-polyacrylamide gel electrophoresis (SDS-PAGE) and analyzed by western blot using anti-Rnasen, anti-Dgcr8 and anti-Dyrk1a antibodies. The arrows indicate protein bands at the expected size. Note that no cross-reaction was found with the lgGs. **(D):** *In situ* proximity ligation assays PLA on primary cortical neurons transfected at DIC5 with Dyrk1a-GFP construct (green fluorescence) and fixed at DIC7, using anti-GFP and anti-Rnasen antibodies (red fluorescence). Non-transfected neurons were used as a negative control. Nuclei were labelled using Topro3 staining (blue fluorescence). Mean interaction point numbers were calculated in nuclear body of at least 25 transfected cortical neurons. *** p<0.0005.

As no *bona fide* antibody was available, we generated a GFP DSCR9 construct, in order to image DSCR9 protein. PLA evidenced nuclear localization of this novel interaction between DSCR9 and CLU. Using in situ PLA, we identified PPI inside nuclei on primary cortical neurons using anti-GFP and anti-Clu antibodies (**Figure 5B**). DSCR9 and DSCR10 have been identified as genes that appeared exclusively in primates such as chimpanzee, gorilla, orangutan, crab-eating monkey and African green monkey and are not present in other non-primate mammals (Takamatsu et al., 2002). CLU gene was identified as one of the 20 LOAD genetic risks (Lambert et al., 2013) and confirmed in a recent meta-analysis a large genome-wide association meta-analysis of clinically diagnosed LOAD (94,437 individuals) (Kunkle et al., 2019). Our results indicate a direct nuclear interaction between the product of a HSA21 gene that contributes to the genomic basis of primate phenotypic uniqueness and a LOAD risk gene.

For DYRK1A-DGRC8 interaction, we first validated this interaction by immunoprecipitation using native conditions (no overexpression) in EK293 cells (**Figure. 5C**). Localization of the interaction in neuronal nuclei was evidenced by PLA on primary cortical neurons transfected at DIC5 with Dyk1a-GFP construct, using anti-GFP and anti-Rnasen antibodies (**Figure. 5D**). The Microprocessor complex is a protein complex involved in the early stages of processing microRNA (miRNA) in animal cells. The complex is minimally composed of the ribonuclease enzyme Drosha and the RNA-binding protein DGCR8 (also known as Pasha) and cleaves primary miRNA substrates to pre-miRNA in the cell nucleus (Wilson and Doudna, 2013).

Deficiency of Dgcr8, a gene disrupted by the 22q11.2 microdeletion that gives a schizophrenia phenotype in humans, results in altered short-term plasticity in the prefrontal cortex (Fénelon et al., 2011). One can suppose that functional phenocopies of this DGC8 haploinsufficiency may occur by titration of its partner RNASEN when DYRK1A is overexpressed. As alterations of the microRNA network cause neurodegenerative disease (Hébert and De Strooper, 2009), our results suggest that DYRK1A-RNASEN interaction could have direct relevance for our understanding of early AD in DS persons.

### Interactome of Hsa21 proteins located in the postsynaptic compartment of the dendritic spine

We next wanted to validate PPI identified by Y2H and to define the subcellular localization of the interactions, using proximity ligation assays. Our working hypothesis was that a subset of Hsa21 proteins and their interactors could be located at the level of dendritic spine. We defined a putative synaptic network (**Figure 6A**) based on the Y2H that we identified and their position in a four layers model as proposed by (Li et al., 2016, 2017). The four layers involve (i) a membrane layer for ionic channels, neurotransmitter receptors and cell-adhesion molecules, (ii) a second layer for DLGs, (iii) a third layer for DLGAPs and (iv) a fourth layer for direct DLGPs interactors (such as SHANKs). Validation of the synaptic network was done using PLA.

**Figure 6.**
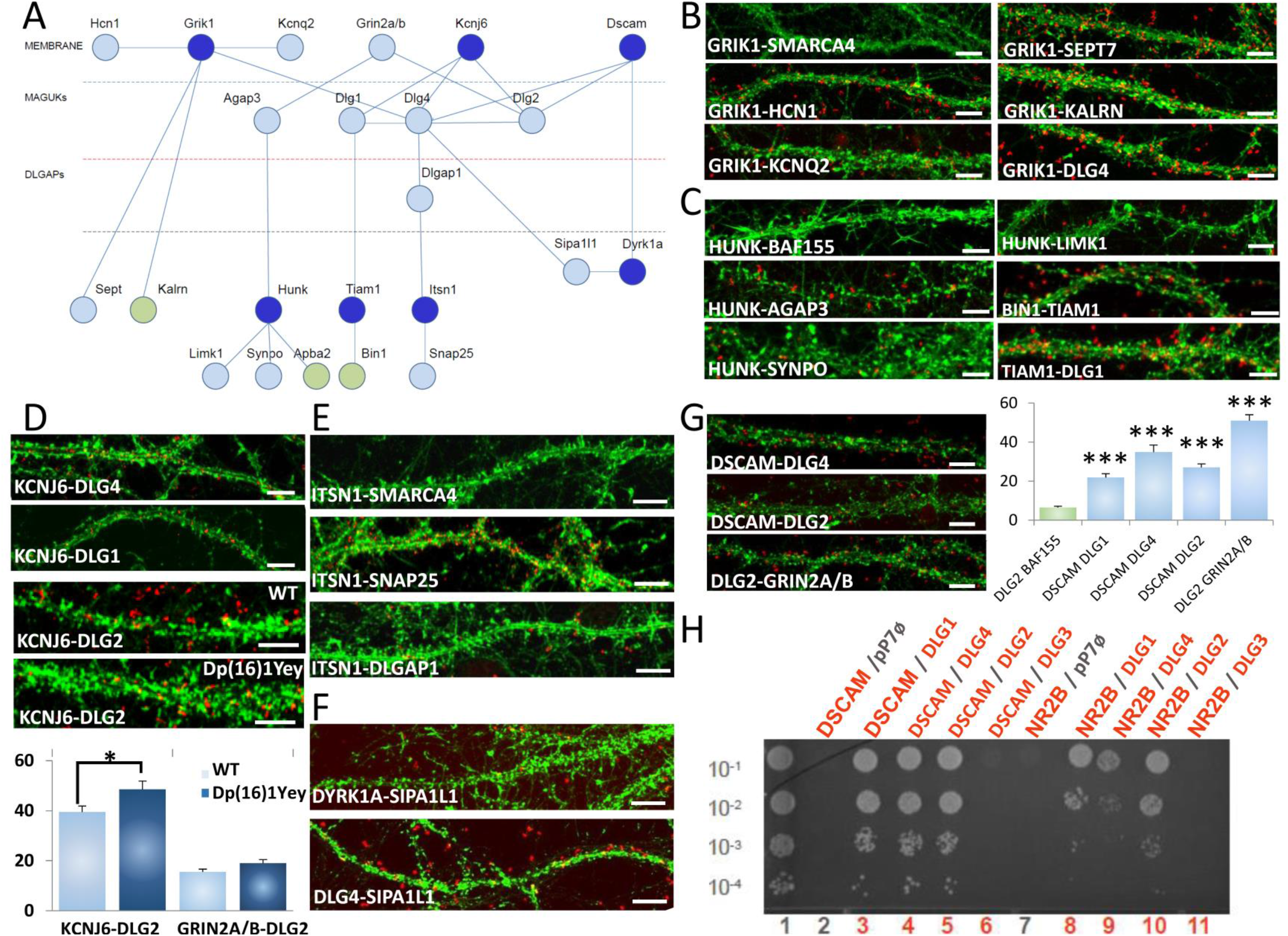
Chr21-encoded proteins have direct interactors in dendritic spine PSD. **A.** Schematic representation of synaptic protein-protein interactions performed by yeast-two-hybrid with the three layers of dendritic spine PSDs. Chr21-encoded proteins and LOAD-related proteins are indicated by dark blue and green circles respectively. **B-G.** *In situ* proximity ligation assays (PLA) on primary cortical neurons fixed at DIC21 (red fluorescence) using anti-Grik1 and anti-Hcn1, anti-Kcnq2, anti-Sept7,anti-Kalrn or anti-Dlg4 antibodies **(B),** anti-Hunk and anti-Agap3, anti-Synpo and anti-Limk1 and also anti-Tiam1 and anti-Bin1 or anti-Dlg1 **(C),** anti-Kcnj6 and anti-Dlg4 or anti-Dlg2 antibodies **(D),** anti-ltsn1 and anti-Snap25 or anti-Dlgap1 **(E),** anti-Sipa1l1 and anti-Dyrk1a or anti-Dlg4 antibodies **(F),** anti-Dscam and anti-Dlg2 or anti-Dlg4 or anti-Grin2a/b antibodies **(G).** PLA using anti-Grik1 and anti-Dlg4 **(B),** anti-Hunk and anti-Synpo **(C),** anti-Kcnj6 and anti-Dlg1 **(D),** anti-ltsn1 and anti-Snap25 **(E),** anti-Sipa1l1 and anti-Dlg4 **(F)** and anti-Dlg2 and anti-Grin2ab **(G)** were performed as positive controls. PLA using either anti-Smarca4 or anti-Baf155 antibodies were performed as negative control. Dendritic network and dendritic spines were labelled using phalloidin staining (green fluorescence). Mean interaction point numbers were calculated in dendrites of 25 to 30 cortical neurons at DIC21. Please see Supplementary Figure for negative controls. d. *In situ* PLA on transgenic Dp(16)1Yey and WT primary cortical neurons fixed at DIC21 (red fluorescence) using anti-Dlg2 and anti-Kcnj6 or anti-Grin2ab antibodies. Dendritic network and dendritic spines were labelled using phalloidin staining (green fluorescence). Mean interaction point numbers were calculated in dendrites of at least 26 cortical neurons at DIC21 (from 3 different embryos per genotype). * p < 0.05 Scale bars=10µm. **H.** Yeast two-hybrid one-by-one assays revealed DSCAM and NR2B as interactors of some of DLGs. Lane 1 is the positive control. Lanes 2 and 7 are the negative controls (pP7-DSCAM or pP7-NR2B vector with empty pP7 vector). Lanes 3 to 6 and 8 to 11 are the DSCAM and NR2B interactions respectively.

We were able to detect 21 PPIs at the level of dendritic spine for which 13 were novel interactions (GRIK1-HCN1; GRIK1-KCNQ2; GRIK1-SEPT7; GRIK1-KALRN; HUNK- AGAP3; HUNK-SYNPO; HUNK-LIMK1; TIAM1-BIN1; TIAM1-DLG1; KCNJ6-DLG4; KCNJ6-DLG2; DSCAM-DLG4; DSCAM-DLG2) and 8 interactions already documented in BioGrid but not validated at the synapse (GRIK1-DLG4; KCNJ6-DLG1; ITSN1-SNAP25; ITSN1-DLGAP1; SIPA1L1-DYRK1A; SIPA1L1-DLG4;DLG2-GRIN2A; DLG2-GRIN2B).

We first focused on protein-protein interactions of Hsa21 GRIK1 (**Figure 6B**). This protein is one of the GRIK subunits functioning as a ligand-gated ion channel. Kainate receptors (KARs) are found ubiquitously in the central nervous system (CNS) and are present pre- and post-synaptically (Lerma and Marques, 2013). We first identified the GRIK1-KCNQ2 interaction. As KCNQ2 potassium channels are known to functionally interact with HCN1 potassium channels in prefrontal cortical dendritic spines (Arnsten et al., 2012), we validated that HCN1, KCNQ2 and GRIK1 physically interact in dendritic shafts and spines using PLA and found a direct interaction between GRIK1 and HCN1 in these compartments. We also identified and validated interactions between GRIK1 with SEPT7 and KALRN. SEPT7, a member of the septin family of GTPases, is localized to dendritic branching points and spine necks (Tada et al., 2007). KALRN is a Rho-GEF exclusively localized to the postsynaptic side of excitatory synapses (Penzes and Jones, 2008) and binds NMDA receptor subunit Nr2b (Grin2b) (Kiraly et al., 2011). These results suggest that GRIK1 is part of two synaptic complexes, one located near PSD-95 (DLG4) at the tip of the dendritic spine and the other at the neck of spines.

Hsa21 HUNK (alias MAK-V) was found in the Y2H screen to interact with the GTPase-activating protein AGAP3, the actin-associated protein Synaptopodin (SYNPO) and the synapse related LIMK1. These three interactions were validated at the level of synapse using PLA (**Figure 6C**). AGAP3 was recently identified as an essential signaling component of the NMDA receptor complex that links NMDA receptor activation to AMPA receptor trafficking (Oku and Huganir, 2013). SYNPO was localized to necks of dendritic spines and linked to the spine apparatus, assumingly playing an essential role in regulating synaptic plasticity (Korkotian et al., 2014). These results suggest that HUNK is involved in complexes localized both near PSD-95 and the spine apparatus. *LIMK1* functions in intracellular signaling and is strongly expressed in the brain, with hemizygosity suggested to impair visuospatial constructive cognition (Frangiskakis et al., 1996).

We next analyzed Hsa21 TIAM1 interaction with BIN1 and DLG1. TIAM1 is a Rac1 associated GEF 1, involved in synaptic plasticity (Penzes and Rafalovich, 2012) and specifically expressed in subgroups of glutamatergic synapses such as dendritic spines of the performant path-dentate gyrus hippocampal synapse (Rao et al., 2019). BIN1 is the second risk factor, after APOE4, identified by Genome Wide Association Studies (GWAS) in LOAD (Kunkle et al., 2019; Lambert et al., 2013). BIN1 protein has multiple functions, including a role at the postsynapse (Daudin et al., 2018; Schürmann et al., 2019). Using PLA, we evidenced TIAM1-BIN1 and TIAM1-DLG1 interactions at the level of dendritic spines (**Figure 6C**).

Another set of noteworthy interactions identified in the Y2H screen (**Figure 6A**), as well as validated via PLA (**Figure 6D**), is between Hsa21 potassium channel KCNJ6, a voltage-insensitive potassium channel member of the kainate ionotropic glutamate receptor (GRIK) family, and three DLG members: DLG1, DLG2 and DLG4. We also found an increase in the number of KCNJ6-DLG2 interactions in the Dp(16)1Yey transgenic mouse model, compared to the control. In contrast, the number of GRIN2A/B-DLG2 interactions was unchanged (**Figure 5D; Supplementary Figure S1**). *KCNJ6* is expressed in dendrites and dendritic spines at the PSD of excitatory synapses (Drake et al., 1997; Luján et al., 2009) and its trisomy induces synaptic and behavioral changes (Cooper et al., 2012). Two interesting interactions detected in the Y2H screen (**Figure 6A**) and validated via PLA (**Figure 6E**) are between Hsa21 Intersectin (ITSN1) and both SNAP25 and DLG- Associated Protein 1 (DLGAP1/GKAP). SNAP25, a member of the SNARE protein family, is not only essential for the exocytosis of synaptic vesicles (Südhof and Rothman, 2009), but involved in trafficking of postsynaptic NMDA receptors (Jurado et al., 2013) and spine morphogenesis (Tomasoni et al., 2013). DLGAP1 is a core protein of the scaffolding complex of the synapse (Kim et al., 1997). These results are in agreement with the *Itsn1* mutant mice phenotype that is characterized by severe deficits in spatial learning and contextual fear memory (Sengar et al., 2013) and in synaptic hippocampal plasticity (Jakob et al., 2017).

SIPA1L1, also known as SPAR is a Rap-specific GTPase-activating protein (RapGAP), a regulator of actin dynamics and dendritic spine morphology, leading to its degradation through the ubiquitin-proteasome system (Pak and Sheng, 2003; Pak et al., 2001). Here, we evidenced direct interaction between SIPA1L1 and DYRK1A or DLG4 (**Figure 6F**) DYRK1A and DLG4

We also identified the location of the Y2H interactions of DSCAM with DLG2 (Discs large 2) and DLG4 at the dendritic spines, using PLA (**Figure 6G**). Then, using DSCAM as bait against single DLG family members (one-by-one Y2H approach), we identified interactions between DSCAM and the four members of the DLG family: DLG1, DLG2, DLG3, and DLG4 (**Figure 6G-H** **Supplementary Figure S2**). DSCAM is known to regulate dendrite arborization and spine formation during cortical circuit development (Maynard and Stein, 2012). DLG1 (alias SAP97), DLG2 (alias PSD93/chapsyn-110) and DLG4 (alias PSD- 95/SAP90) are known to bind various proteins and signaling molecules at the postsynaptic density (PSD) (Kim and Sheng, 2004; Sheng and Kim, 2011). Intriguingly, mice lacking *Dlg2* and people with *Dlg2* mutations display abnormal cognitive abilities (Nithianantharajah et al., 2013).

Altogether, our results demonstrate location of novel proteins in the nanoscale region that corresponds to the four layers of the dendritic spine and covers a 75 nm-thick region from synaptic cleft (Tao et al., 2018).

### DSCAM-DYRK1A interaction

In our Y2H screens, we identified interactions between human DSCAM, its human paralog DSCAML1 and human DYRK1A. This interaction occurs in the same intracellular domain (45 amino acids) of these cell adhesion molecules (**Figure 7A**). Furthermore, their drosophila homologs, DSCAM4 and minibrain (MNB), were also found to interact in the same phylogenetically conserved domain of 45 amino acids (**Figure 7B**). We confirmed human DSCAM-DYRK1A interaction by immunoprecipitation, using adult mouse cortex (**Figure 7C**). We were unable to find an anti DYRK1A antibody that can be used for PLA proximity ligation assay. We used preparation of synaptosomes and immunoprecipation to evidence DSCAM-DYRK1A interaction in this subcellular fraction (**Figure 7D-E**). We found that DSCAM interacts with DLG4 and DLG2 in the postsynapse, using PLA (**Figure 6G**). Altogether, our data indicate that DSCAM-DYRK1A and DLGs interacts in the postsynapse. To the best of our knowledge, this is the first localization of DSCAM and DYRK1A in the postsynapse.

**Figure 7.**
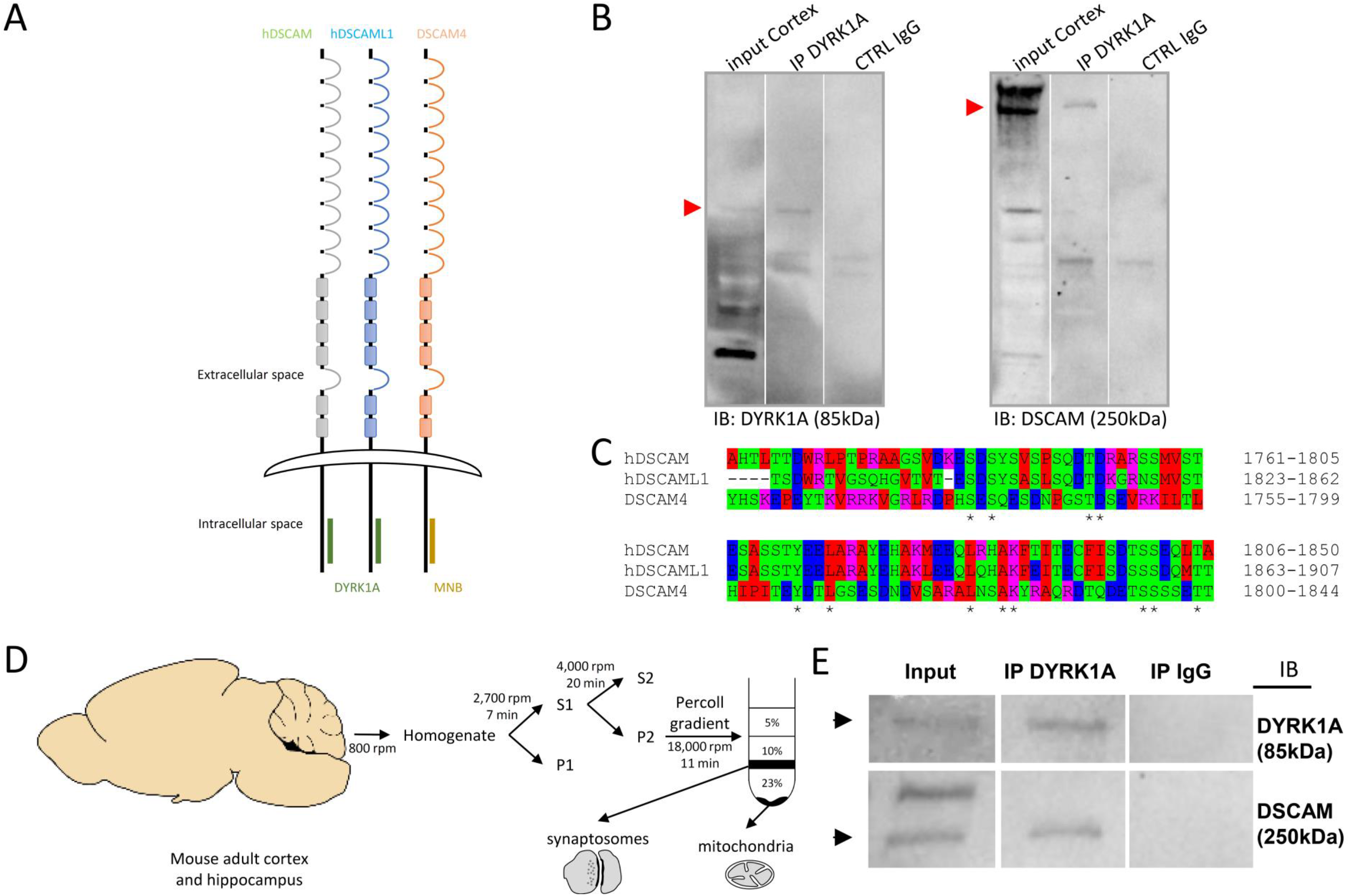
Conservation of DSCAM-DYRK1A interaction through evolution. **A.** Schematic representation of DSCAM and DYRK1A protein family interaction. Human DSCAM (hDSCAM in green), human DSCAML1 (hDSCAML1 in blue) and its drosophila orthologus (dDSCAM4 in red) share the same conserved proteic domain interacting with human DYRK1A (hDYRK1A) or its drosophila orthologus (MNB) respectively. **B.** Adult mouse cortical protein extract were immunoprecipitated (IP) using anti-Dyrk1a antibody and anti-lgG antiboby as a negative control. The input and precipitated fractions were then resolved by sodium dodecyl sulphate-polyacrylamide gel electrophoresis (SDS-PAGE) and analyzed by western blot using anti-Dyrk1a and anti-Dscam antibody. Red arrows indicate protein bands at the expected size. Note that no cross-reaction was found with the lgGs. **C.** Amino acid alignment of the DSCAM domain that interacts with DYRK1A and Minibrain. This alignment was performed with ClustalW 2.1 software. **D.** Schematic representation of synaptosome enrichment protocol. **E.** Adult mouse cortical protein extracts were immunoprecipitated (IP) using anti-Dyrk1a antibody and anti-lgG antibody as a negative control. The input and precipitated fractions were then resolved by sodium dodecyl sulphate-polyacrylamide gel electrophoresis (SDS-PAGE) and analyzed by western blot using anti-Dyrk1a and anti-Dscam antibody. Note the band of 85 kDa expected for the Dyrk1a protein and the 250 kDa band expected for Dscam protein. No cross-reaction was found with the lgGs.

**Figure 8.**
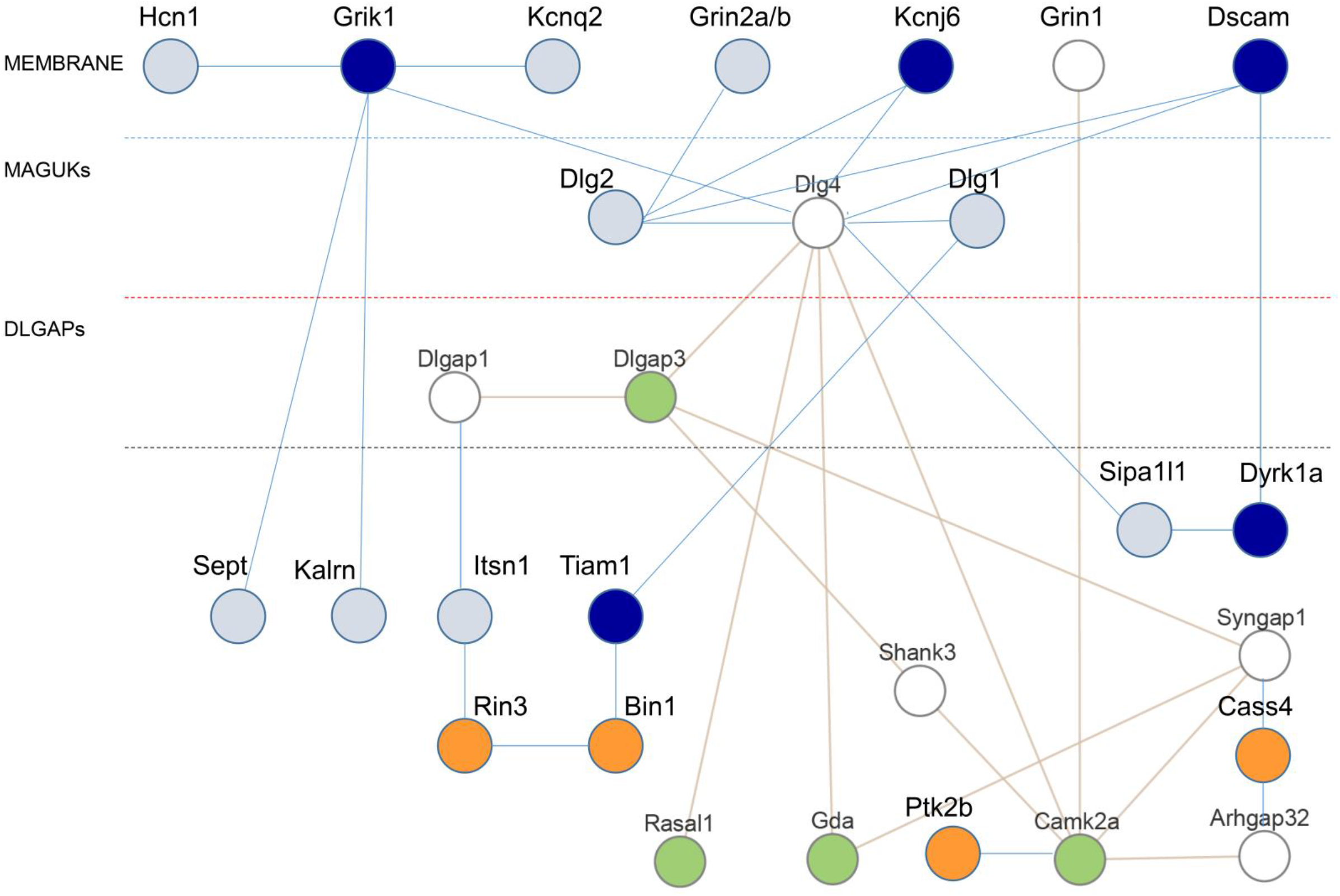
Protein-protein interactions in the three layers of dendritic spine PSD: enrichment in proteins encoded by either HSA21 or LOAD-GWAS genes. Schematic representation of synaptic protein-protein interactions performed by yeast-two-hybrid, with the three layers of dendritic spine PSDs indicated (membrane; MAGUKs, DLGAPs). HSA21-encoded proteins are represented as dark blue circles. LOAD-GWAS encoded proteins are represented by dark orange circles.

Both DSCAM and DYRK1A are part of a subset of 30 genes that confers 20-fold risk in autism-spectrum disorders (ASDs). In ASD, studies leveraging the statistical power afforded by rare de novo putatively damaging variants have identified more than 65 strongly associated genes (Sanders et al., 2015). The most deleterious variants (likely gene disrupting or LGD) in the highest confidence subset of these genes (N = 30) as a group confer approximately 20-fold increases in risk, with LGD variants in the highest confidence genes carrying even greater risks (Willsey et al., 2018).

### A synaptic network enriched in Hsa21 proteins, ASD high-risk genes, ARC-related protein network and Late Onset Alzheimer Diseases (LOAD) risk factors

We summarized our PPI data at the synapse with a protein network of 28 products (**Figure 7**). This synaptic network is enriched in HSA21 proteins with 5 out 234 HSA21 gene products (the results are enriched 18.72 fold compared to expectations; hypergeometric p-value = 6.435 e- 07). This 28 proteins network is also enriched in high-risk ASDs genes. We found that DSCAM and DYRK1A are part of this network that includes also DLGAP1 and SHANK3. These four genes are considered to confer ∼20-fold increases in risk, for a group containing 30 genes (Sanders et al., 2015; Willsey et al., 2018). The enrichment here is 97.35 fold compared to expectations (hypergeometric p-value = 7.52e-08; parameters: 4, 28, 30, 20444). [234/Ensembl release 88 - Mar 2017]

The third group of enrichment is an ARC-dependent postsynaptic complex involved with Neural Dysfunction and Intelligence (Fernández et al., 2017). This group includes 20 proteins and 7 of them (Grin2a, Grin1, Dlg4, Dlg2, Dlgap1, Syngap1, Camk2a) are part of our synaptic complex (the results are enriched 255.55 fold compared to expectations; hypergeometric p-value = 3.06e-16; parameters: 7, 28, 20, 20444). The fourth group is Late-onset (LOAD) that involve 11 new loci corresponding to 26 candidate genes, (Karch et al., 2014; Lambert et al., 2013). We found an enriched 112.33 fold compared to expectations (hypergeometric p-value = 4.12e-08; parameters: 4, 28, 26, 20444).

Altogether, these results demonstrate that our postsynaptic network is enriched in Hsa21 proteins, ASD high-risk genes, ARC-related protein network and LOAD risk factors.

## Discussion

In spite of the availability of various mouse DS models, cognitive impairment phenotypes found in DS have not been related to specific alterations of molecular pathways. Furthermore, to the best of our knowledge, no specific pathways linked to synaptic alterations have been described in models overexpressing a given chr21 gene as compared to models overexpressing a syntenic region.

In the present study, we analyzed molecular changes in the hippocampus of several mouse DS models. We found molecular changes linked to chromatin remodeling in the Dyrk1A BAC 189N3 mice. In contrast, expression of an extra copy of the entire Hsa21 syntenic region, spanning 22.9 Mb and containing 115 Hsa21 gene orthologs, including *Dyrk1A*, on Mmu16 in the Dp(16)1Yey transgenic mouse model, induced changes in GO glutamatergic and gabaergic synaptic transmission.

Using large scale Y2H we evidenced that both direct interactors (n=1687) and their second order interactors that we capture using our rebound screen (n=1949) are enriched in ID genes. This observation suggests that protein-protein complexes that includes a Hsa21 protein are at-risk complexes for ID.

The PLA approach that allows localizing Protein-Protein Interactions in a given subcellular comportment allows us to focus on synaptic compartment as a potential candidate where subtle deregulations may occur (Grant, 2018; Koopmans et al., 2019).

We identified that interactions at the synapse are enriched in Hsa21 gene products. In particular, we were able to demonstrate that both DYRK1A and DSCAM are part of the postsynapse. DYRK1A and DSCAM are high-risk genes for ASDs with a 20 fold increase as compared to the general population (Sanders et al., 2015; Willsey et al., 2018). Gene dosage changes of both DYRK1A and DSCAM may involve distinct molecular mechanisms as compared to pathways changes occurring in ASDs and linked to damaging mutations involving either DYRK1A or DSCAM. Further work to decipher molecular changes in respective pathways can illuminate both DS and ASDs pathophysiology.

We also characterized a significant enrichment in ARC-dependent synaptic network involved in intelligence and brain diseases (Fernández et al., 2017). Mutations disrupting this molecular hierarchy change the architecture of synaptome maps, potentially accounting for the behavioral phenotypes associated with neurological and psychiatric disorders (Grant, 2018). Further work will be needed to analyze the changes in protein complexes at the synapse, when gen dosage change for partners of these complexes.

It is well documented that Down syndrome, caused by trisomy of chromosome 21, is the single most common risk factor for early-onset Alzheimer’s disease. Triplication of APP has been as considered as a candidate for this phenotype but trisomy of human chromosome 21 enhances amyloid-β deposition independently of an extra copy of APP indicating that triplication of chromosome 21 genes other than APP is likely to have an important role to play in Alzheimer’s disease pathogenesis in individuals who have Down syndrome (Wiseman et al., 2015b). We reported here an enrichment in LOAD genes that have been identified by GWAS strategies, with RIN3, BIN1 and CASS4 in our postsynaptic network. We (Daudin et al., 2018) and other (Schürmann et al., 2019) characterized BIN1 at the level of the synapse using super-resolution microscopy. Other protein network that includes BIN1 and CASS4 has been reported in microglia (Nott et al., 2019). Altogether, these results suggest that cell-distinct proteins complex may contribute to Alzheimer phenotype in DS.

Interestingly, from our Y2H and PLA approaches, we identified novel candidates in the postsynaptic domain that we characterized by four layers as in (Li et al., 2016, 2017) has a very restricted width in the range of 75 nm (Tao et al., 2018). Super-resolution microscopy approaches recently revealed that spine synapses *in vitro* and brain slices nanodomains that form a trans-synaptic column and contain discrete, precisely aligned sub-diffraction nanomodules, whose number, not size, scales with spine volume (Hruska et al., 2018; Tang et al., 2016).

Changes in stoichiometry of interactors, as expected for Hsa21 proteins may modify the functional impact of a given protein complex. The report that a same Neuroligin4 mutation can generate either ID or high-level ASD supports such subtle changes (Laumonnier et al., 2004). Similarly, some protein complexes may integrate only a given form of a protein, as it is the case for TIAM1 in glutamatergic synapses from entorhinal cortex (Rao et al., 2019).

Together, our results suggest that protein-protein interactions identified here can be part of different dendritic spine signalosomes deregulated by three doses of Hsa21 proteins.

In conclusion, our results provide the first report, to our knowledge, of differential impacts of chromosome 21 *DYRK1A* on chromatin remodeling and of the 115 Hsa21 gene orthologs including *DYRK1A*, on synapse function. Our results exemplify the link of DS to other forms of ID and to degenerative diseases displaying complex genetics such as LOAD. The molecular pathways studied here can promote the development of novel therapeutic targets to treat cognitive impairments found in DS.

## MATERIALS AND METHODS

### Animals and genotyping

All experiments were approved by the Institut National de la Santé et de la Recherche Médicale (INSERM) animal care D-13-055-19 agreement (to V.C.), 03882.02 and B751403 agreements (to M.S.), in agreement with the European community council directive 2010/63/UE.

We used wild-type mice of the OF1 strain for neuronal primary culture, wild-type of the C57BL6 strain and Tg(Dyrk1a)189N3Yah (named 189N3) or Dp(16Lipi-Zfp295)1Yey (named Dp(16)1Yey) transgenic lines for neuronal primary cultures. Genotypes were determined using genomic DNA extracted from skeletal muscle fragments.

### RNA Sequencing

#### Sample preparation

The hippocampi were dissected from genotyped E16-E18 embryos (n= 3 or 4 per genotype for 189N3 or Dp(16)1Yey transgenic mouse, respectively). Sample were homogenized in Trizol reagent (GIBCO), purified on nucleospin column (Macherey Nagel), treated with DNase I (Ambion) and processed according to the manufacturer’s instructions.

#### Sequencing

The RNA sequencing was performed on Illumina HiSeq Sequencers (from the High Throughput Platform, CNG, CEA, Evry, France).

#### Analysis

The Bowtie-TopHat-Cufflinks pipeline was used as previously described (Trapnell C et al., 2012). Reads were mapped on *Mus musculus* mm10 genome and the UCSC known genes was used as transcriptome index. Cuffmerges were run on all samples. The merged assembly were mapped on the Gencode (release M4) main annotation and all the transcripts which were not described within were removed (antisens genes, unknown transcript) in order to focus on protein coding genes. For the quantification (cuffquant) of the abundance, the frag-bias-correct and the multi-read-correct options of the program on the merged assembly were used. The differential analysis was performed on two levels: gene level and transcript isoform level.

### Primary cell cultures

Primary cultures from OF1 mice were performed as described in Loe-Mie et al., 2010. Heterozygous 189N3 or Dp(16)1Yey mice were crossed with C57BL6, resulting in embryos of transgenic or wild type genotypes. E15.5 189N3 or Dp(16)1Yey cortical neurons were dissociated by individually dissecting each embryo out of its amniotic sac, removing the head and dissecting out the target brain tissue in an separate dish. The remainder of the brain was used for genotyping. Neurons from each embryo were dissociated enzymatically (0.25% trypsin), mechanically triturated with a flamed Pasteur pipette, and individually plated on 24- well dishes (1×10^5^ cells per well) coated with poly-DL-ornithine (Sigma), in DMEM (Invitrogen) supplemented with 10% fetal bovine serum. Four hours after plating, DMEM was replaced by Neurobasal® medium (Invitrogen) supplemented with 2mM glutamine and 2% B27 (Invitrogen). For nuclear interactions or dendritic interactions, cortical neurons were analyzed after 7 days or 21 days in culture respectively.

### Constructs

Mouse Dyrk1a cDNA was cloned in GFP plasmid as described (Lepagnol-Betel et al. 2009). Human USP25 and SYK were cloned in GFP and MYC plasmids respectively as described (Cholay et al. 2010). Human GDI1 and DSCR9 were amplified by PCR from IMAGE: 4156714 and IMAGE: 6065320 cDNA clones, respectively (SourceBioscienes) with the following primers:

> GDI1 forward: 5’-gatcggccggacgggccGACGAGGAATACGATGATCGTG

> GDI1 reverse: 3’-gatcggccccagtggccTCACTGCTCAGCTTCTCCAAAGACGTC

> DSCR9 forward: 5’-gatcggccggacgggccATGGGCAGGATTTGCCCCGTGAAC

> DSCR9 reverse: 3’-gatcggccccagtggccTCACCATAATTCCTGTGTCTGAATCTGAA

The SfI digestion products of the amplicons were inserted into the multiple cloning site of the HA and GFP expression vectors respectively under control of the CMV promoter.

### Primary cell cultures and transfection

Cortical primary neurons were cultured as described above. At DIC5, the cells were transfected with constructs using Lipofectamine 2000 (Invitrogen), as described by the manufacturer. Cells were analyzed 48 hours after transfection at DIC7.

### HEK293 cell cultures and transfection

HEK293 cell line were plated in 24-well plates in DMEM (Invitrogen) supplemented with 10% fetal bovine serum. At 70% confluency, the cells were transfected with constructs (co-transfections were performed at 1:3 ratio) using Lipofectamine 2000 (Invitrogen), as described by the manufacturer. Cells were analyzed 48 hours after transfection.

### *In situ* proximity ligation assays (PLA) and microscopy

Cells were fixed by incubation for 20 min at room temperature in 4% paraformaldehyde in phosphate-buffered saline (PBS), permeabilized by incubation for 10 min at room temperature in 0.3% Triton X-100 in PBS, washed two times within PBS and PLA was realized according to the instructions of the manufacturer (DuoLink, Sigma). Primary antibodies used were as shown in Supplementary Table S4. For the analysis of PLA interactions points, cells were scanned using the laser scanning confocal microscope (Leica, SP5 from PICPEN imagery platform Centre de Psychiatrie et Neuroscience) at 63× magnification, and Z-stacks were build using the ImageJ software (Wayne Rasband, NIH). Nuclear PLA interaction number was manually counted inside the nuclear body and normalized with the nuclear area of each neuron. Synaptic PLA interaction number was manually counted on 150 μm long dendritic segments starting after the first branch point in the dendritic tree. One 150 μm dendritic segment per neuron was analyzed.

### Statistical analysis

The analyses performed on transgenic neurons with at least 3 embryos and at least 10 cells per embryo for synaptic and nuclear analyses. The analyses performed on OF1 neurons with at least 3 different cultures and at 8 cells and 14 cells per culture for synaptic and nuclear analyses respectively. The analyses performed on HEK293 cells with at least 3 different transfections and 5 cells per transfection for nuclear or cytoplasmic analyses. T-tests were performed with Excel Software.

### Protein extraction and Western blot analysis

HEK293 cells or mouse cortex (pool from three adult OF1 mice) were homogenized on ice in Tris-buffered saline (100 mM NaCl, 20 mM Tris-HCl, pH 7.4, 1% NP40, 1 x CIP). The homogenates were centrifuged at 13,000g for 10 min at 4°C and the supernatants were stored at -80 °C. Cell lysate protein concentration was determined using the BCA Protein assay kit (Thermofisher). For SDS-PAGE, 40μg of protein was diluted in Laemmli 1x (BioRad) with DTT and incubate for 5mn at 95°C. Proteic samples were loaded in each lane of a 4-15% precast polyacrylamide gel (BioRad) and ran in Mini-Protean at 200V in Tris/Glycine running buffer (BioRad). Following SDS-PAGE, proteins were semi-dry electroblotted onto nitrocellulose membranes using the Trans-Blot Turbo Transfer System (BioRad). Membranes were incubated for 1 h at room temperature in blocking solution (PBS 1x containing 5% non-fat dried milk, 0.05% Tween 20) and then for overnight at 4°C with the primary antibody. Primary antibodies used were as shown in Supplementary Table S4. Membranes were washed in PBS 1x containing 0.05% tween 20 and incubated for 1 h at room temperature with anti-mouse, anti-rabbit or anti-goat HRP-conjugated secondary antibody. Membranes were washed three times in PBS 1x containing 0.05% tween 20. Immune complexes were visualized using the Clarity Western ECL Substrate (BioRad). Chemiluminescence was detected using the ChemiDoc XRS Imaging System (BioRad). As secondary antibodies, we used protein A or protein G IgG, HRP-conjugated whole antibody (1/5,000; Abcam ab7460 or ab7456 respectively).

### Immunoprecipitation

1mg of protein extracts were incubated, after preclear with 50µL of dynabeads (Novex), 3h at 4°C under rotating with 10 µg of primary antibody (Supplementary Table S4; anti-mouse and anti-rabbit whole IgG (Millipore 12-371 and 12-370 respectively)). Add 50µL of protein A or protein G dynabeads and incubate 30mn at 4°C under rotating. Protein-antibody complexes were washed four times in 100 mM NaCl, 20 mM Tris-HCl, pH 7.4, 1% NP40 and analyzed by immunoblot.

### Laser-assisted microdissection, Total RNA preparation and quantitative real-time PCR (Q-RT-PCR) analysis

Embryonic left and right hippocampus was microdissected from genotyped P21 mouse brains using a laser-assisted capture microscope (Leica ASLMD instrument) with Leica polyethylene naphthalate membrane slides as described in Lepagnol-Bestel et al, 2009. RNA preparation and Q-RT-PCR are performed as described in Lepagnol-Bestel et al, 2009. Q-RT-PCR results are expressed in arbitrary unit.

#### Statistical analysis

All data are shown as means+/-SEM. Statistics were performed using IgorPro (Wavemetrics), and statistical significance was determined by the Student’s t test (two tailed distribution, paired) unless otherwise stated.

#### Reagents

Stock solutions were prepared in water or DMSO, depending on the manufacturers’ recommendation, and stored at -20°C. Upon experimentation, reagents were bath applied following dilution into ACSF (1/1000). D-APV and DCGIV were purchased from Tocris Bioscience. Salts for making cutting solution and ACSF were purchased from Sigma.

### Networks bioinformatics analyses

We used Disease Association Protein-Protein Link Evaluator (DAPPLE) that looks for significant physical connectivity among proteins encoded for by genes in loci associated to disease (Rossin et al., 2011). Interactions are extracted from the database “InWeb” that high confidence interactions. Connections can be direct and indirect. The significance of the interaction parameters are tested using a permutation method that compares the original network with thousands of networks created by randomly re-assigning the protein names while keeping the overall structure (size and number of interactions) of the original network.

To complement the DAPPLE analysis, we used the WebGestalt suite (Liao et al (Liao et al., 2019).

### Yeast-two-hybrid experiments

#### Orthologs collection, multiple sequence alignments, and phylogenetic trees

A list of 234 genes from Hsa21 was examined. For each gene, orthologs across mammals were determined.

### Y2H library

We used a human Adult brain poly(A^+^) RNA (Invitrogen: Discovery Line Human normal Brain mRNA, Sex: M, Age: 27, Catalog No: D6030-15, LOT No: A308079) constructed in the pP6 plasmid derived from the original pACT222 and transformed in *E. coli* (DH10B; Invitrogen, Carlsbad, CA, USA). The complexity of the primary libraries was over 50 million clones. Sequence analysis was performed on 300 randomly chosen clones to establish the general characteristics of each library. The libraries were then transformed into yeast by classical lithium acetate protocol. Ten million independent yeast colonies were collected, pooled and stored at −80°C as equivalent aliquot fractions of the same library (Fromont-Racine et al., 1997; Fromont-Racine et al., 2002).

Two-hybrid screens were performed using a cell to cell mating protocol. For each bait, a test screen was performed to adapt the screening condition. The selectivity of the HIS3 reporter gene was eventually modulated with 3-aminotriazole (Sigma, St Louis, MO, USA) in order to obtain a maximum of 285 histidine-positive clones for 50 millions diploids screened. For all the selected clones, lacZ activity was estimated by overlay assay on solid media in 96-well plate format. Inserts of all positive clones were amplified by PCR22, 23 and then sequenced on an ABI 3700 automatic sequencer (Applied Biosystem, Foster City, CA, USA).

### Prey identification

5′ and 3′ sequences were determined for all positive clones in a screen. These were in turn filtered for quality using PHRED and ALU repeats were masked. Sequence contigs were built using CAP324 and searched against the latest release of GenBank using BLASTN

### Identifying reliable interactions

Interactions were filtered based on a <u>p</u>redicted <u>b</u>iological <u>s</u>core (PBS) (Fromont-Racine et al., 2002) The PBS was calculated based on randomly sequenced cDNA library and adopts the conventional form of a *P*-value, where the smaller the PBS (*P*-value) the more significant. The PBS relies on two different levels of analysis. Firstly, a local score takes into account the redundancy and independency of prey fragments, as well as the distribution of reading frames and stop codons in overlapping fragments. Secondly, a global score takes into account the interactions found in all the screens performed at Hybrigenics using the same library. This global score represents the probability of an interaction being nonspecific. For practical use, the scores were divided into four categories, from A (highest confidence) to D (lowest confidence). A fifth category (E) specifically flags interactions involving highly connected prey domains previously found several times in screens performed on libraries derived from the same organism. Finally, several of these highly connected domains have been confirmed as false-positives of the technique and are now tagged as F. The PBS scores have been shown to positively correlate with the biological significance of interactions .

#### Preparation of bait constructs

The coding sequence for each bait protein was PCR-amplified and cloned in-frame with the LexA DNA binding domain (DBD) into plasmid pB27, derived from the original pBTM116. DBD constructs were checked by sequencing the entire insert. Several inserts were cloned in-frame with the Gal4 DBD into plasmid pB66, derived from combines data from a variety of public PPI sources including MINT, BIND, IntAct and KEGG and defines pAS2ΔΔ (24). For DSCR8 (contested coding gene) cDNA coding for MKEPGPNFVTVRKGLHSFKMAFVKHLLLFLSPRLECSGSITDHCSLHLPVQEILMSQPPEQLGLQTNLGNQESSGMMKLFMPRPKVLAQYESIQFMP have been used.

#### Preparation of Dyrk1a ΔpolyHis mutant bait construct

The sequence coding for Dyrk1a C-terminus (aa 600-763) was modified to remove the poly-Histidine stretch to prevent potential artefacts of binding with Cysteine-rich prey proteins without altering the folding of the bait. This region was modify by genesynthesis (Eurofins-Genomics) inserted in the cDNA cloned resulting in the sequence behind: DYRK1A delta polyHis: PQQNALHAAHGNSSAAAGAHAGAAHAHGQQALGNRTRP

#### Preparation of prey constructs 1-by1 assays

DLG1 (aa 305-653), DLG4 (154-356), DLG2- 5 (aa 84-440) prey plasmids were extracted from the diploid cells obtained from the Y2H screening with wild-type DSCAM of the Human Adult Brain library. Inserts are cloned in-frame with the Gal4 activation domain (AD) into plasmid pP6, derived from the original pGADGH. The coding sequence for DLG3 (aa 212-385) was PCR-amplified and cloned in-frame with the Gal4 AD into plasmid pP7. The AD constructs were checked by sequencing.

#### Y2H Screening and 1by1 interaction assays

Bait and prey constructs were transformed in the yeast haploid cells, respectively CG1945 or L40ΔGal4 (mata) and YHGX13 (Y187 ade2- 101::loxP-kanMX-loxP, matα) strains. The diploid yeast cells were obtained using a mating protocol with both yeast strains (Fromont-Racine et al., 2002) His+ colonies were selected on a medium lacking tryptophan, leucine and histidine, and supplemented with 3-aminotriazole to handle bait autoactivation when necessary. The prey fragments of the positive clones were amplified by PCR and sequenced at their 5’ and 3’ junctions. The resulting sequences were used to identify the corresponding interacting proteins in the GenBank database (NCBI) using a fully automated procedure.

Interaction pairs were tested in duplicate as two independent clones from each diploid were picked for the growth assay. For each interaction, several dilutions (10-1, 10-2, 10-3 and 10-4) of the diploid yeast cells (culture normalized at 5×104 cells) and expressing both bait and prey constructs were spotted on several selective media. The DO-2 selective medium lacking tryptophan and leucine was used as a growth control and to verify the presence of both the bait and prey plasmids. The different dilutions were also spotted on a selective medium without tryptophan, leucine and histidine (DO-3). Four different concentrations of 3-AT, an inhibitor of the HIS3 gene product, were added to the DO-3 plates to increase stringency. The following 3-AT concentrations were tested: 1, 5, 10 and 50 mM.

## ACCESSION CODES

Data are available at Intact, EBI-EMBL under the accession number IM-27626.

## ACKNOWLEDGMENTS

We would like to acknowledge the funding support from the European Union’s Seventh Framework Programme for research, technological development and demonstration AgedBrainSYSBIO under grant agreement No. 305299 (to M.S.) and Fondation Lejeune. This work was also partly funded by INSERM, European JPND (TransPathND) and CNES (to M.S.).

## Notes

### Competing Interest Statement

The authors have declared no competing interest.

### Summary of Updates

We added RNAseq paragraph in Methods

